# *cis-* and *trans*-regulatory factors contributing to divergent activity of the *TDH3* promoter in *Saccharomyces* yeast

**DOI:** 10.64898/2026.04.01.715911

**Authors:** Mohammad A. Siddiq, Hannah P. Kania, Nicholas J. Brown, Patricia J. Wittkopp

**Author notes:** **Corresponding author:** Patricia J. Wittkopp.

## Abstract

Changes in regulatory sequences controlling the timing and activity of gene products underlie much of natural phenotypic variation. Yet, what these changes are and how they impact gene expression remain largely unknown. To address this question, we investigated how transcriptional activity and homeostatic responsiveness of orthologous promoters of the metabolic gene *TDH3* evolved among *Saccharomyces* yeast. We found that promoter expression level increased specifically in the *S. cerevisiae* lineage and that a substantial part of this increase was caused by genetic variants located between the well-characterized, conserved binding sites for two direct transcriptional regulators. These nucleotide changes altered the promoters’ expression levels while leaving the expression dynamics conserved. Further, the effects of these nucleotide changes were only seen in the presence of a third transcription factor, TYE7p, which is recruited by the other transcription factors through protein-protein interactions. These results suggest that the *cis*-regulatory changes act through their influence on the collective assembly/activation of the transcription factors, and that changes acting through such a mechanism can allow distinct parts of gene expression, such as expression level and dynamics, to be tuned separately.

## Main Text

Molecular processes that regulate the expression of genes in response to developmental and environmental cues are critical for determining the relationship between genetic and phenotypic variation. Genetic variants that affect gene regulatory mechanisms have become increasingly appreciated as the cause of phenotypic variation among populations and between species. This evolutionary importance of variation in gene regulation was presciently hypothesized 50 years ago (Britten and Davidson 1969; King and Wilson 1975), and investigations of phenotypic variation at various levels, from metabolism to morphology, have strengthened this hypothesis in the decades since (Shubin et al. 1997; Stern and Orgogozo 2008; Blount et al. 2012; Martin and Orgogozo 2013; Coyle and King 2025). Understanding the molecular and evolutionary processes that cause regulatory sequences to be the tools of evolutionary tinkering and the mechanisms that govern the evolution of regulatory sequences remains a pressing challenge for the field.

Expression of each gene is controlled by interactions between *cis*- and *trans*-regulatory factors, both of which contribute to the evolution of gene expression (Hill et al. 2021). *cis*-regulatory factors include DNA sequences (e.g., promoters, enhancers, UTRs) that affect the expression at a particular locus, and *trans*-regulatory factors include diffusible molecules (e.g., transcription factors, regulatory RNAs) present in a cell which often regulate the expression of multiple genes from many loci. Despite progress in understanding the function of *cis*-regulatory sequences and predicting the *trans*-acting transcription factors that interact with them (de Boer et al. 2020; Avsec et al. 2021; Vaishnav et al. 2022; Mahendrawada et al. 2025; Xie et al. 2025), our ability to predict which genetic changes contribute to an evolutionary change in gene expression remains poor (Nora et al. 2023). Most often, mutations in transcription factor binding sites are assumed to be responsible for *cis*-regulatory divergence (Wray 2007), perhaps because mutations in transcription factor binding sites tend to have large effects on gene expression. Mutations with large effects, however, might not be the most likely to fix during evolution (Rockman 2012; Umans et al. 2021). To better understand the mechanisms of *cis*-regulatory divergence, specific nucleotide changes and the molecular causes of their contribution to differences in *cis-*regulatory activity between species need to be identified for more genes.

In the baker’s yeast *Saccharomyces cerevisiae*, the *TDH3* promoter has been used as a model system for understanding how genetic changes in both *cis*- and *trans*-acting sequences can introduce variation in gene expression and how selection shapes that variation within species (reviewed in (Wittkopp 2023)). The *TDH3* promoter controls transcription of the most highly expressed of the three paralogous genes in *S. cerevisiae* encoding the core metabolic protein glyceraldehyde-3-phosphate dehydrogenase (McAlister and Holland 1985). It is regulated by four transcription factors: Rap1p, Gcr1p, Gcr2p, and Tye7p (Uemura and Jigami 1992; Deminoff and Santangelo 2001; Holland et al. 2019; Shively et al. 2019; Bergenholm et al. 2021). Rap1p and Gcr1p cooperatively recognize sequence elements in the promoter (Tornow et al. 1993; Deminoff and Santangelo 2001), Gcr2p forms a heteromeric complex with Gcr1p (Uemura and Jigami 1992; Deminoff and Santangelo 2001), and Tye7p is thought to be recruited through protein-protein interactions with the other transcription factors (Holland et al. 2019; Shively et al. 2019). Quantitative variation in *TDH3* activity directly affects fitness during fermentative and respirative growth (Duveau, Toubiana, et al. 2017; Siddiq et al. 2024). Further, the activity of the *S. cerevisiae TDH3* promoter is dynamically sensitive to internal and external cellular conditions: promoter expression changes based on glucose availability during culture growth, and loss of *TDH3* activity triggers a homeostatic feedback mechanism mediated by upregulation of Gcr1p (Vande Zande et al. 2023). Studies of mutations and polymorphisms in *S. cerevisiae* have demonstrated that the activity of the *TDH3* promoter is strongly constrained, with mutations to the RAP1p and GCR1p binding sites being particularly costly (Metzger et al. 2015; Metzger et al. 2016; Duveau, Yuan, et al. 2017; Duveau, Toubiana, et al. 2017; Duveau et al. 2018; Metzger and Wittkopp 2019; Duveau et al. 2021; Vande Zande et al. 2022; Siddiq et al. 2024). Yet, despite this constraint, comparative RNA-seq analyses show that the activity level of the *TDH3* promoter has diverged through *cis-*regulatory differences between *S. cerevisiae* and its sister species *S. paradoxus* (Krieger et al. 2020). We therefore sought to uncover the genetic changes through which the orthologous *TDH3* promoters’ activity diverged and, in turn, better understand the mechanistic causes of evolution of gene expression.

The genetic changes responsible for the divergent activity of the *TDH3* promoter between *S. cerevisiae* and *S. paradoxus* arose since the species last shared a common ancestor, approximately 5-10 million years ago (Replansky et al. 2008). To infer the most likely direction of evolutionary change between these two species, we examined the activity of the *TDH3* promoter from *Saccharomyces mikatae* and *Saccharomyces kudriavzevii*, which last shared a common ancestor with *S. cerevisiae* and *S. paradoxus* 10-15 and 15-20 million years ago, respectively (Replansky et al. 2008). We used each species’ promoter to drive expression of a yellow fluorescent protein (YFP) in a reporter gene that was integrated at the *HO* locus of the *S. cerevisiae* genome (**Figure 1A**). These orthologous promoters ranged from 668 to 678 bp long and ranged from 73% to 85% sequence identity (**Figure 1B**). YFP fluorescence was used as a proxy for *TDH3* promoter activity, and fluorescence was measured using flow cytometry in at least 45,000 cells in each of 3 replicate populations for each of the four reporter genes in populations of cells grown in rich media (YPD). Cultures were grown at 30°C, and yeast cells were sampled at the end of the fermentative growth stage (∼18-21 hours). After correcting each individual cell for its estimated cell size, we took the median fluorescence of each sample, averaged these medians among the replicates of the same strain, and compared the expression levels driven by the orthologous promoters using a linear mixed model with a random effect to account for day-to-day variation. We found that the *S. cerevisiae TDH3* promoter drove a significantly higher level of expression than the other three species, which all drove similar levels of expression (**Figure 1C; Supplementary Table 1A**).

**Figure 1.**
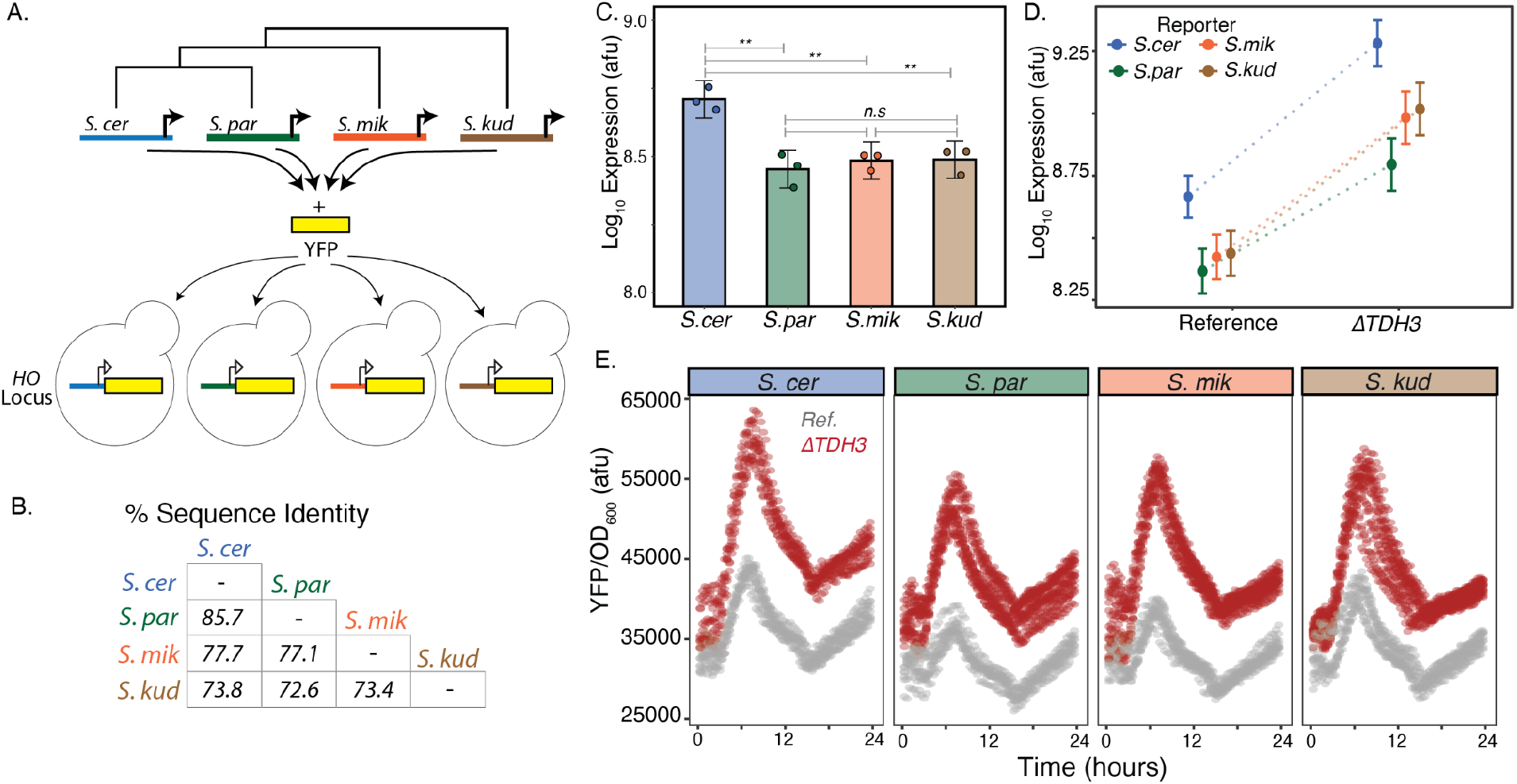
Divergent activity of the *TDH3* promoter activity in *S. cerevisiae*. (A) Orthologous *TDH3* promoters from *S. cerevisiae* (*S. cer*), *S. paradoxus* (*S. par*), *S. mikatae* (*S. mik*), and *S. kudriavzevii* (*S. kud*) were fused to the *venusYFP* coding sequence (YFP) to make a series of reporter genes. Each reporter gene was inserted into the *HO* locus of a common *S. cerevisiae* reference strain. (B) Pairwise nucleotide sequence identity is shown for the different species’ *TDH3* promoters. (C) Mean activity of orthologous promoters as determined by flow cytometry (YFP expression, normalized for cell size) is shown with error bars indicating the standard error of the mean. Dots show the mean values for individual replicates, each consisting of >40,000 cells and collected on a different day. Asterisks designate statistically significant differences among genotypes, as determined using a linear model to estimate the effects of each genotype with the Tukey HSD method for post-hoc pairwise comparisons (*S. cer* vs. *S. par*: *t* = 5.27, *p* = 0.0003; *S. cer* vs. *S. mik*: *t* = 4.27, *p* = 0.002; *S. cer* vs. *S. kud*: *t* = 4.01, *p* = 0.004; all other comparisons had *p* > 0.5). (D) Activity of the orthologous *TDH3* promoter alleles in genomic backgrounds with (Reference) and without a functional *TDH3* gene (*ΔTDH3*); the different promoters were upregulated similarly in response to the deletion of *TDH3* (*F* = 1.1988; p =0.3394). Points and error bars show estimated means and standard errors, respectively. (E) YFP fluorescence driven by *TDH3* promoter alleles in the reference (gray) and *TDH3* mutant (red) backgrounds during growth for 24 hours on liquid YPD. Data is shown for 4-8 replicates of each of the 8 genotypes.

To determine whether the sequence divergence among these promoters also affected the homeostatic feedback mechanism reported for the *TDH3* promoter in *S. cerevisiae* (Vande Zande et al. 2023), we used the same procedure to measure expression driven by each promoter in a strain of *S. cerevisiae* with the native *TDH3* gene deleted (TDH3::ΔTDH3). In all cases, we found that deletion of the *TDH3* gene increased the activity of the *TDH3* promoter, showing that this compensatory response was conserved among species (**Figure 1D**). This conserved dynamic behavior of the *TDH3* promoter was not confined to homeostatic regulation -- we also measured activity of the species-specific *TDH3* promoter alleles in *S. cerevisiae*, with and without a functional *TDH3* gene, over 24 hours of growth when cells are undergoing a diauxic shift on YPD and found that that all four promoters, in both genetic backgrounds, showed a similar pattern of expression change over time, even though the *S. cerevisiae* promoter retained the highest overall expression (**Figure 1E; Supplementary Table 1B**). These data show that derived sequence changes in the *S. cerevisiae TDH3* promoter increased the promoter’s activity level without altering its dynamic homeostatic and metabolic regulation. This finding is consistent with genomic comparisons of *S. cerevisiae* and *S. paradoxus* showing that expression level often evolves independently of expression dynamics (Krieger et al. 2020; Shih and Fay 2021) and that divergence in expression level tends to be caused by *cis*-acting genetic changes.

To try to identify specific nucleotide changes contributing to higher activity of the *S. cerevisiae TDH3* promoter, we focused on a region containing an upstream activation sequence responsible for *TDH3* expression (Bitter et al. 1991). We first looked for changes in experimentally validated binding sites that recruit the Rap1p (**R**epressor-**a**ctivator **p**rotein) and the heteromer of Gcr1p and Gcr2p (**G**ly**c**olysis **r**egulator 1 and 2; hereafter Gcr1p/2p) transcription factors (Bitter et al. 1991; Chambers et al. 1995; Metzger et al. 2015). These proteins cooperatively activate the transcription of *TDH3* as well as other glycolysis genes (Mizuno et al. 2004). We found that these transcription factor binding sites were highly conserved in sequence and position among all four orthologous promoters (**Figure 2A**), suggesting that they are not the source of the divergent activity in *S. cerevisiae*. However, we noticed several differences in the 14 base-pair region between the Gcr1p/2p and Rap1p binding sites (**Figure 2A**). Comparing the full *S. cerevisiae* and *S. paradoxus TDH3* promoter sequences showed that this region was more than twice as divergent (5/14 bp, 36%) as the rest of the promoter (92/664 bp, 14%; Fisher’s exact test; p = 0.037). We suspected these changes may be functionally consequential due to their physical proximity to the binding motifs, potentially modifying the efficacy of TF-DNA interactions.

**Figure 2.**
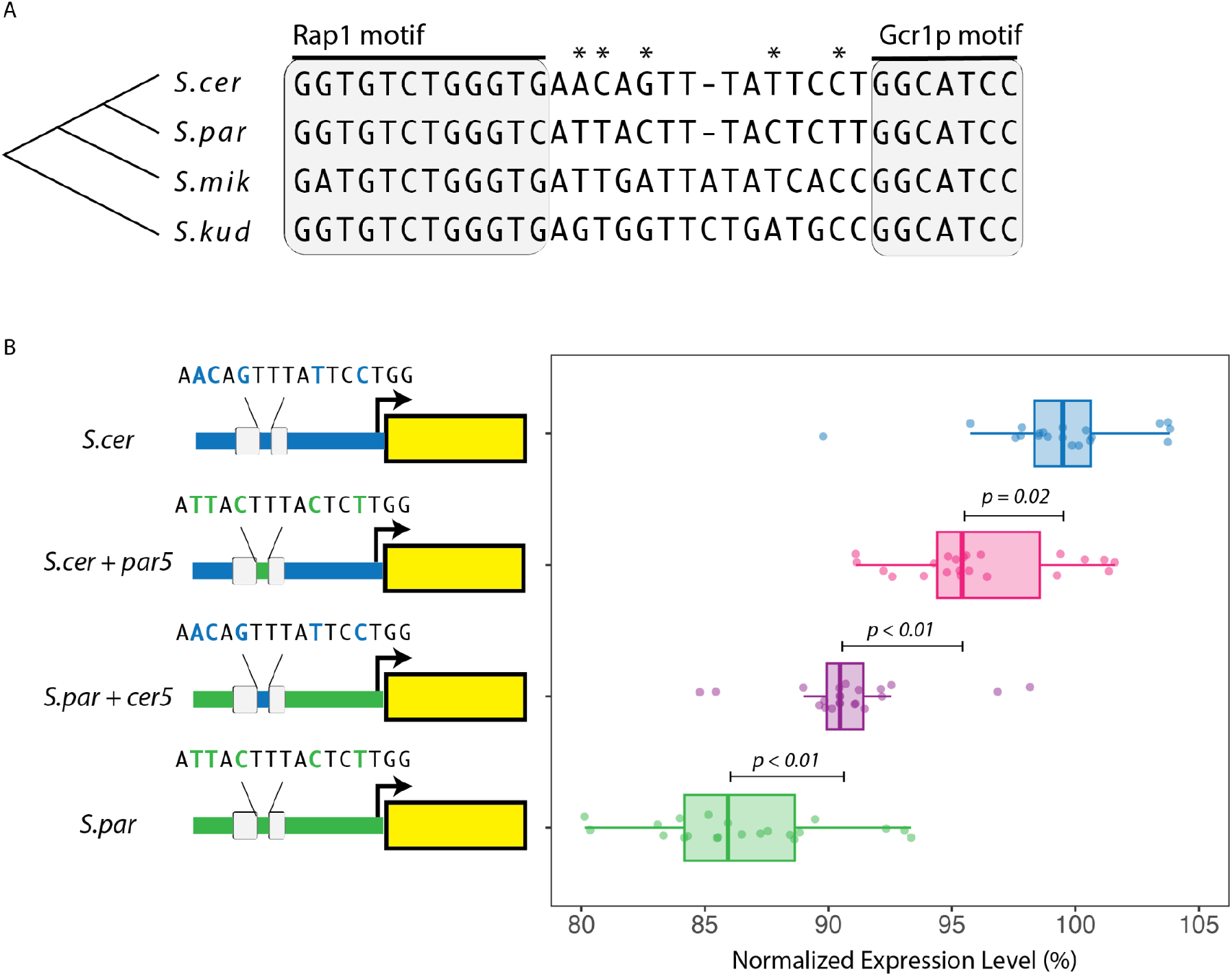
Sequence divergence between the Rap1p and Gcr1p binding sites affects activity of the *TDH3* promoter. (A) Sequence alignment shows a region of the *TDH3* promoter from *S. cerevisiae* (*S. cer*), *S. paradoxus* (*S. par*), *S. mikatae* (*S. mik*), and *S. kudriavzevii* (*S. kud*) containing a previously characterized upstream activating sequence. This sequence includes binding sites for the transcription factors *Rap1p* and *Gcr1p*. High-affinity binding sites for both of these transcriptional regulators, shown in gray, are highly conserved across species. Asterisks highlight five positions between these binding sites that differentiate the *S. cerevisiae* allele from the *S. paradoxus* allele. (B) Relative activity of the *S. cerevisiae* (*S. cer*) and *S. paradoxus* (*S. par*) *TDH3* promoter alleles are shown alongside activity of recombinant constructs in which the five divergent sites indicated with asterisks in panel A were swapped between the species-specific promoters (*S*.*cer + par5* and *S. par + cer5*), as shown in the schematics. Each data point plotted represents a separate experimental replicate (>20,000 cells/measurement). *P-*values displayed for pairwise differences were calculated from a linear model fit to the data with the Tukey HSD method for post-hoc comparisons.

We tested whether these changes contribute to divergence of the *TDH3* promoter by swapping the 5 divergent sites in this 14 bp region between the *S. cerevisiae* and *S. paradoxus TDH3* promoters and assaying activity of these chimeric promoters using the same YFP reporter gene inserted at the HO locus (**Figure 2B**). We found that these 5 nucleotides did indeed have a significant effect on expression of the reporter genes: the *S. paradoxus* promoter with the *S. cerevisiae* alleles had higher expression than the wild-type *S. paradoxus* promoter (**Figure 2B**; t = 3.97, p < 0.001, Tukey HSD), and the *S. cerevisiae* promoter with the *S. paradoxus* alleles had lower expression than the wild-type *S. cerevisiae* promoter (**Figure 2B**, t = 2.91, p = 0.02, Tukey HSD). In neither case, however, was changing these 5 nucleotides sufficient to fully convert the expression level from one species to the other. Rather, our data show that changing these 5 nucleotides is sufficient to explain ∼35% of the expression level divergence between the two promoters. The rest of the divergence in expression level between *S. cerevisiae* and *S. paradoxus* is presumably caused by one or more of the 87 other divergent sites between the promoters of these two species located outside of this region.

The Rap1p and Gcr1p/2p transcription factors binding to sites that define the ends of this region form a complex and cooperatively regulate *TDH3* expression (Mizuno et al. 2024). Consequently, sequence divergence in this intervening region might affect the formation and/or activity of this complex. The Tye7p transcription factor, a basic helix-loop-helix (bHLH) protein that binds to E-box motifs (Gordan et al. 2013), also regulates *TDH3* expression as a part of this complex (Holland et al. 2019, Liu et al. 2020, Bergenholm et al. 2021) (**Figure 3A**). However, despite having a DNA recognition helix, Tye7p cannot directly bind the *TDH3* promoter or other glycolytic promoters on its own (Shively et al. 2019). Instead, it is recruited to the promoter through interactions with Gcr2p in the Gcr1p/2p heteromer, after which it can make direct contacts with DNA as a part of this complex (Liu et al. 2020). Consistent with this observation, the 14 bp region does not contain any matches (strictly or more relaxed) to the E-Box binding motif (CACGTG). Based on this information, we hypothesized that 5 sequence differences between the Rap1p and Gcr1p/2p binding sites might alter activity of the *TDH3* promoter by altering the formation or activity of this complex -- perhaps through changing the interactions of Rap1p or Gcr1p/2p with Tye7p.

**Figure 3.**
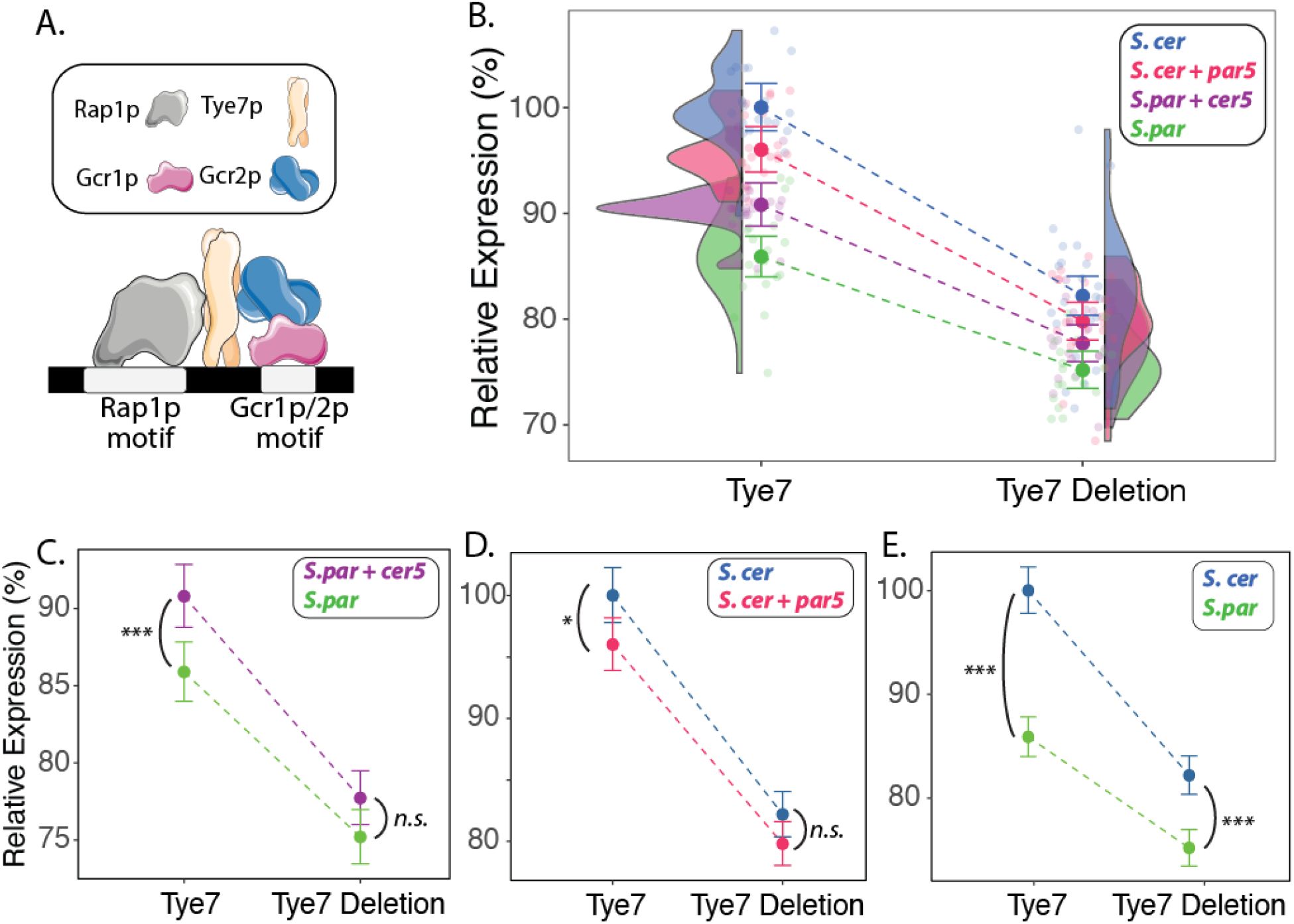
Effects of divergent sites depend on Tye7p. (A) Schematic shows a model of Tye7p recruitment by the Gcr1p/2p complex and Rap1p, as described in Shively et al. (2019). Binding motifs for Gcr1p and Rap1p are necessary for proper *TDH3* promoter activation; Tye7p requires no specific binding motif to localize to the *TDH3* promoter. However, we hypothesize that certain DNA sequences at the site of Tye7p localization might better stabilize the collective complex and lead to greater transcriptional activation. (B) The relative activity of the *S. cerevisiae* (*S*.*cer*), *S. paradoxus* (*S*.*par*), and two chimeric *TDH3* promoter alleles (*S*.*cer + par5* and *S. par + cer5*) is shown in the presence (Tye7) and absence (Tye7 Deletion) of Tye7p. Opaque points and error bars display estimated mean values for each genotype with 95% confidence intervals; transparent points represent observations from individual replicates. Density distributions are also shown, describing all replicates for each promoter allele. (C-E) Same data as B but highlighting the Tye7-dependent effects of changing the 5 nucleotides between the Rap1p and Gcr1p binding sites in the *S. paradoxus TDH3* promoter (C), the *S. cerevisiae TDH3* promoter (D), as well as the overall effect of the combined 92 differences between the *S. cerevisiae* and *S. paradoxus* promoters (E). Statistical significance of pairwise comparisons between promoters is shown (values in **Supplementary Table 2**). P-values were calculated from a linear model with reporter, genetic background, and random effect of day with the Tukey HSD method. N.S. = *p > 0*.*10;* * = *p < 0*.*05* ; ** = *p < 0*.*01* ; *** = *p < 0*.*05*.

To test whether the effects of the 5 differences were dependent on Tye7, we compared the expression of the YFP reporter genes driven by the *S. cerevisiae, S. paradoxus*, and two chimeric *TDH3* promoters in a genomic background in which the *TYE7* gene had been deleted (TYE7::ΔTYE7). After measuring YFP fluorescence for each of these 8 strains, we used a linear model to estimate the individual and interaction effects of the promoter and *TYE7* genotypes, accounting for day-to-day variation as a random effect. We found that all 4 promoters displayed lower activity when measured in the TYE7::ΔTYE7 strain (**Figure 3B**; *F* = 563.3, *p* < 0.001, Type III ANOVA), indicating that Tye7p is required for normal *TDH3* promoter activity in both *S. cerevisiae* and *S. paradoxus*. However, there was a significant interaction between promoter and *TYE7* genotype (*F* = 4.06, *p* < 0.01, Type III ANOVA), indicating that removing Tye7p had different effects on different promoters. Specifically, in absence of Tye7p, the *S. cerevisiae* alleles of the 5 divergent sites no longer caused a significant increase in expression level when introduced into the *S. paradoxus TDH3* promoter (**Figure 3C;** t = 2.29, p = 0.10, Tukey HSD). Similarly, introducing the *S. paradoxus* alleles for these 5 sites into the *S. cerevisiae* promoter no longer led to a significant decrease in expression (**Figure 3D;** t = 2.11, p = 0.16, Tukey HSD). These data show that the effects of the 5 divergent sites between the Rap1p and Gcr1p/2p binding sites are mediated by regulatory interactions involving Tye7p. Consistent with these findings, the difference in activity between the *S. cerevisiae* and *S. paradoxus TDH3* promoters decreased by 49% in the absence of Tye7p, though the remaining difference was still statistically significant (**Figure 3E;** t = 6.18, p < 0.001).

This study reveals an important, and perhaps underappreciated, mechanism by which *cis*-regulatory sequences evolve: mutations outside of canonical binding sites for transcription factors can drive substantial changes in gene expression by modulating protein-protein interactions within regulatory complexes. While binding sites for the direct regulators of *TDH3* (Rap1p and Gcr1p) remain conserved between *S. cerevisiae* and *S. paradoxus*, nucleotide changes in the intervening sequence have increased *TDH3* promoter activity in *S. cerevisiae* by altering regulation mediated by Tye7p, which cannot bind DNA alone at this locus. These changes did not create or abolish a binding motif for Tye7p. Instead, they were likely able to produce this effect because of the mechanism through which Tye7p activates expression as part of a transcription factor collective (Junion et al. 2012; Liu et al. 2020). The initial recruitment of Tye7p and relative strength of activation by Tye7p happen in a stepwise way through protein-protein and protein-DNA interactions, respectively, in which specific Tye7-DNA interactions at the site of recruitment can quantitatively alter expression level even though they are not necessary for complex formation (Liu et al. 2020). As such, changes in these interactions can allow variation in expression level to evolve without altering the upstream regulators and signals to which the promoter responds, enabling different aspects of gene expression to evolve separately.

This finding is counter to the prevailing view that genetic variation in transcription factor binding sites is the primary driver of *cis*-regulatory evolution; rather, our findings highlight the importance of sequences that may influence low-affinity interactions, cooperativity, and assembly of multi-protein regulatory complexes. These factors, increasingly appreciated in molecular and developmental studies of gene regulation (Kribelbauer et al. 2019), are likely to provide powerful frameworks for explaining how variation in gene regulation evolves. As we expand our catalogs of evolutionarily significant regulatory mutations to include more genes and species, a more nuanced view of *cis*-regulatory evolution will likely emerge -- one in which changes that fine-tune protein complex formation, rather than simply disrupting or creating transcription factor binding sites, play a central role in shaping gene expression and ultimately contributing to phenotypic diversity.

## Materials and Methods

### Yeast Strains

The strains of *S. cerevisiae* used in this study were all derived from the standard S288c strain and had an alpha mating type. These strains had previously been modified to carry alleles of *RM1, TAO3, CAT5* and *MIP1* that increase sporulation efficiency and decrease petite frequency relative to the native alleles of the S288c, as described in Metzger et al. (Metzger et al. 2016). To quantify promoter expression, a reference non-fluorescent strain was engineered to carry different promoter-YFP constructs at a common position in the *HO* locus. The full cassette included the *TDH3* promoter allele, a Venus YFP reporter, a Cyc1 terminator, and a KanMX resistance cassette. To swap out the different promoter alleles, an intermediate strain was created at the *HO* locus carrying the KanMX marker, a target site for CRISPR-Cas9 but lacking a promoter-YFP reporter construct; the different promoter-reporter constructs, including the reference *S. cerevisiae TDH3* promoter allele, were then precisely introduced into this background using the same CRISPR-Cas9 target site using a modified version of the pML104 plasmid and accompanying protocol described in Laughery et al. (Laughery et al. 2015). The open-reading frames for *TDH3* and *TYE7* were also removed using a similar CRISPR-Cas9 based strategy, albeit with 20bp targets that were specific to the different loci (ACACACATAAACAAACAAAA for *TDH3*; TAGTCATATCAACGTCAACA for *Tye7*). For all strains generated using CRISPR-Cas9, the transformed cells were plated on SC-uracil and grown for 2-3 days at 30ºC. Colonies were screened for the correct genotype at the loci of interest with Sanger sequencing, and colonies with the desired changes were subsequently cured of the pML104 plasmid by selection on 5FOA media. Finally, the strains were grown in liquid YPD (10 g/L yeast extract, 20 g/L peptone, 20 g/L dextrose) until saturation, mixed with glycerol for a final concentration of 20% glycerol, and stored at -80ºC.

Annotated sequences of the constructs used at the HO locus are provided in **Supplementary File 1**. Annotated sequences of the reference and deletion variants at the *TDH3* and *Tye7* locus, as well as a summary of yeast strains, are provided in **Supplementary File 2**.

### Plate Reader Assays

Strains with different genotypes were patched from glycerol stocks onto YPG agar media (10 g/L yeast extract, 20 g/L peptone, 20 g/L agar, 20 g/L glycerol) and grown at 30ºC for 2–3 days. Colonies were subsequently picked, placed onto 96-well 2 mL deep-well plates (randomized across wells) in 1 mL of liquid YPD, and grown with shaking for approximately 48 hours at 30ºC. At this point, the strains had acclimated to growth on glucose and reached saturation. The strains were then diluted into 96-well plates, with 5 µl of saturated culture added to 195 µl of YPD. The plates were then incubated at 30ºC with shaking in a BioTek Synergy H1 plate reader (Agilent), with readings of OD660 and YFP fluorescence taken at 20-minute intervals for 48 hours. Each genotype was included in at least three separate wells per replicate, and three replicates were measured for each genotype.

### Flow Cytometry Experiments

Strains with different genotypes were patched from glycerol stocks onto YPG agar media and grown at 30ºC for 2–3 days and subsequently transferred to YPD agar media to acclimate to growth on glucose. Colonies were then picked and grown in 1 mL of liquid YPD at 30ºC on a shaker or a rotating wheel for approximately ∼17-21 hours, which is the stage when growth slows down following the exhaustion of sugar and prior to the diauxic shift into aerobic metabolism. Cells at this stage were diluted to an OD660 between 0.5 and 0.8 in 1x PBS, and these samples were analyzed by flow cytometry using an Attune NXT4 flow cytometer. Forward Scatter height, width, and area (FSC.H, FSC.W, and FSC.A, respectively) were used to identify populations of singlets, and the 488 nm excitation laser and 530/30 emission filter were used to record their associated fluorescent values.

Cell size is positively correlated with fluorescence, and we adjusted for differences in cell size using a principal components analysis in R using the *flowClust (Lo et al. 2009)* and *flowCore* (Hahne et al. 2009) packages. We took the population of cells that fit our criteria for single cells (e.g, no evidence of cell division or more than one cell in a droplet), estimated the logarithm of the forward scatter (FSC.A) and fluorescence intensity (BL1.A), defined the vector (*v*) between the origin and intersection of the two eigenvectors, and calculated the angle (θ) between the first eigenvector and *v*. This rotation angle reflects the residual correlation between size and fluorescence that persists after linear scaling; we therefore transformed FSC.A and BL1.A by a rotation of angle θ centered on the intersection of the eigenvectors, then divided the transformed FL1.A by the transformed FSC.A to obtain cell-size-corrected fluorescence values. These corrected values were used for downstream statistical analyses.

### Statistical analyses

We used mixed-effect linear models to estimate the effect of different genetic factors on expression levels and account for random day-to-day variation in R using the *lme4 (Bates et al. 2015), lmerTest* (Kuznetsova et al. 2017), and *emmeans* (Lenth and Piaskowski 2026) packages. To estimate the effects of the reporter genotype and the presence/absence of *TDH3* in the genetic background, we fit the following linear model: ***Fluorescence ∼ Reporter * TDH3 Genotype + (1***|***Date)***. We used the *emmeans* package to estimate the marginal means and contrasts among Reporter and *TDH3* Genotype levels. Contrasts with Tukey adjusted p-values < 0.05 were deemed to be statistically significant.

We used a similar framework for testing for the effects of the 5 nucleotides that varied between *S. paradoxus* and *S. cerevisiae* promoters as well as their dependence on Tye7p. For these experiments, we had three technical replicates for each genotype on each day (biological replicate). We used this replicate data to first estimate the mean fluorescence level of our reference genotype -- the unmutated *S. cerevisiae TDH3* reporter construct in an unmutated background-- and scaled the fluorescence level of each sample by this number. We then calculated the mean scaled fluorescence level for each reporter genotype for each day and removed background autofluorescence, which was estimated using a fluorescence-null strain. The mixed effect linear model used to analyze these data was specified as ***Scaled Fluorescence ∼ Reporter * Tye7 Genotype + (1***|***Date)***. We estimated the marginal means and statistically significant differences among Reporter and Tye7 genotypes as described above.

## Acknowledgments

We thank Anna Redhuis, Erick Bayala, and all other members of the Wittkopp Laboratory, as well as Professor Ken Cadigan, for valuable discussions of this work. We also thank Professor Laura Buttitta and the University of Michigan Molecular, Cellular, and Developmental Biology core facilities for access to equipment and assistance with flow cytometry.

## Funding

This work was supported by the National Institutes of Health (5F32CA261115 and T32HG000040 to M.A.S. and 5R35GM118073 to P.J.W.) as well as the National Science Foundation (DEB-1911322 and MCB-1929737 to P.J.W.) In addition, M.A.S. was supported by the Michigan Pioneer Fellows program, and N.J.B was supported by the EEB Summer Research and Travel Award and Program in Biology Director’s Award from the University of Michigan. The content is solely the responsibility of the authors and does not necessarily represent the official views of the National Institutes of Health or the National Science Foundation.

## Notes

### Competing Interest Statement

The authors have declared no competing interest.

## References

Avsec Ž, Weilert M, Shrikumar A, Krueger S, Alexandari A, Dalal K, Fropf R, McAnany C, Gagneur J, Kundaje A, et al. 2021. Base-resolution models of transcription-factor binding reveal soft motif syntax. Nat. Genet. 53:354–366.

Bates D, Mächler M, Bolker B, Walker S. 2015. Fitting linear mixed-effects models using lme4. J. Stat. Softw. 67:1–48.

Bergenholm D, Dabirian Y, Ferreira R, Siewers V, David F, Nielsen J. 2021. Rational gRNA design based on transcription factor binding data. Synth. Biol. 6:ysab014.

Bitter GA, Chang KK, Egan KM. 1991. A multi-component upstream activation sequence of the Saccharomyces cerevisiae glyceraldehyde-3-phosphate dehydrogenase gene promoter. Mol. Gen. Genet. 231:22–32.

Blount ZD, Barrick JE, Davidson CJ, Lenski RE. 2012. Genomic analysis of a key innovation in an experimental Escherichia coli population. Nature 489:513–518.

de Boer CG, Vaishnav ED, Sadeh R, Abeyta EL, Friedman N, Regev A. 2020. Deciphering eukaryotic gene-regulatory logic with 100 million random promoters. Nat. Biotechnol. 38:56–65.

Britten RJ, Davidson EH. 1969. Gene regulation for higher cells: a theory. Science 165:349–357.

Chambers A, Packham EA, Graham IR. 1995. Control of glycolytic gene expression in the budding yeast (Saccharomyces cerevisiae). Curr. Genet. 29:1–9.

Coyle MC, King N. 2025. The evolutionary foundations of transcriptional regulation in animals. Nat. Rev. Genet. 26:812–827.

Deminoff SJ, Santangelo GM. 2001. Rap1p requires Gcr1p and Gcr2p homodimers to activate ribosomal protein and glycolytic genes, respectively. Genetics 158:133–143.

Duveau F, Hodgins-Davis A, Metzger BP, Yang B, Tryban S, Walker EA, Lybrook T, Wittkopp PJ. 2018. Fitness effects of altering gene expression noise in Saccharomyces cerevisiae. Elife 7:e37272.

Duveau F, Toubiana W, Wittkopp PJ. 2017. Fitness Effects of Cis-Regulatory Variants in the Saccharomyces cerevisiae TDH3 Promoter. Mol. Biol. Evol. 34:2908–2912.

Duveau F, Vande Zande P, Metzger BP, Diaz CJ, Walker EA, Tryban S, Siddiq MA, Yang B, Wittkopp PJ. 2021. Mutational sources of trans-regulatory variation affecting gene expression in Saccharomyces cerevisiae. eLife 10:e67806

Duveau F, Yuan DC, Metzger BPH, Hodgins-Davis A, Wittkopp PJ. 2017. Effects of mutation and selection on plasticity of a promoter activity in Saccharomyces cerevisiae. Proc. Natl. Acad. Sci. U. S. A. 114:E11218–E11227.

Hahne F, LeMeur N, Brinkman RR, Ellis B, Haaland P, Sarkar D, Spidlen J, Strain E, Gentleman R. 2009. flowCore: a Bioconductor package for high throughput flow cytometry. BMC Bioinformatics 10:1–8.

Hill MS, Vande Zande P, Wittkopp PJ. 2021. Molecular and evolutionary processes generating variation in gene expression. Nat. Rev. Genet. 22:203–215.

Holland P, Bergenholm D, Börlin CS, Liu G, Nielsen J. 2019. Predictive models of eukaryotic transcriptional regulation reveals changes in transcription factor roles and promoter usage between metabolic conditions. Nucleic Acids Res. 47:4986–5000.

Junion G, Spivakov M, Girardot C, Braun M, Gustafson EH, Birney E, Furlong EEM. 2012. A transcription factor collective defines cardiac cell fate and reflects lineage history. Cell 148:473–486.

King MC, Wilson AC. 1975. Evolution at two levels in humans and chimpanzees. Science 188:107–116.

Kribelbauer JF, Rastogi C, Bussemaker HJ, Mann RS. 2019. Low-affinity binding sites and the transcription factor specificity paradox in eukaryotes. Annu. Rev. Cell Dev. Biol. 35:357–379.

Krieger G, Lupo O, Levy AA, Barkai N. 2020. Independent evolution of transcript abundance and gene regulatory dynamics. Genome Res. 30:1000–1011.

Kuznetsova A, Brockhoff PB, Christensen RHB. 2017. LmerTest package: Tests in linear mixed effects models. J. Stat. Softw. 82:1–26.

Laughery MF, Hunter T, Brown A, Hoopes J, Ostbye T, Shumaker T, Wyrick JJ. 2015. New vectors for simple and streamlined CRISPR-Cas9 genome editing in Saccharomyces cerevisiae: Vectors for simple CRISPR-Cas9 genome editing in yeast. Yeast 32:711–720.

Lenth R, Piaskowski J. 2026. emmeans: Estimated Marginal Means, aka Least-Squares Means. R package version 2.0.2, https://rvlenth.github.io/emmeans/.

Liu J, Shively CA, Mitra RD. 2020. Quantitative analysis of transcription factor binding and expression using calling cards reporter arrays. Nucleic Acids Res. 48:e50.

Lo K, Hahne F, Brinkman RR, Gottardo R. 2009. flowClust: a Bioconductor package for automated gating of flow cytometry data. BMC Bioinformatics 10:1–8.

Mahendrawada L, Warfield L, Donczew R, Hahn S. 2025. Low overlap of transcription factor DNA binding and regulatory targets. Nature:1–9.

Martin A, Orgogozo V. 2013. The Loci of repeated evolution: a catalog of genetic hotspots of phenotypic variation. Evolution 67:1235–1250.

McAlister L, Holland MJ. 1985. Isolation and characterization of yeast strains carrying mutations in the glyceraldehyde-3-phosphate dehydrogenase genes. J. Biol. Chem. 260:15013–15018.

Metzger BPH, Duveau F, Yuan DC, Tryban S, Yang B, Wittkopp PJ. 2016. Contrasting Frequencies and Effects of cis- and trans-Regulatory Mutations Affecting Gene Expression. Mol. Biol. Evol. 33:1131–1146.

Metzger BPH, Wittkopp PJ. 2019. Compensatory trans-regulatory alleles minimizing variation in TDH3 expression are common within Saccharomyces cerevisiae. Evol Lett 3:448–461.

Metzger BPH, Yuan DC, Gruber JD, Duveau F, Wittkopp PJ. 2015. Selection on noise constrains variation in a eukaryotic promoter. Nature 521:344–347.

Mizuno T, Kishimoto T, Shinzato T, Haw R, Chambers A, Wood J, Sinclair D, Uemura H. 2004. Role of the N-terminal region of Rap1p in the transcriptional activation of glycolytic genes in Saccharomyces cerevisiae. Yeast 21:851–866.

Nora EP, Aerts S, Wittkopp PJ, Bussemaker HJ, Bulyk M, Sinha S, Zeitlinger J, Crocker J, Fuxman Bass JI. 2023. Emerging questions in transcriptional regulation. Cell Syst. 14:247–251.

Replansky T, Koufopanou V, Greig D, Bell G. 2008. Saccharomyces sensu stricto as a model system for evolution and ecology. Trends Ecol. Evol. 23:494–501.

Rockman MV. 2012. The QTN program and the alleles that matter for evolution: all that’s gold does not glitter. Evolution 66:1–17.

Shih C-H, Fay J. 2021. Cis-regulatory variants affect gene expression dynamics in yeast. Elife 10:e68469.

Shively CA, Liu J, Chen X, Loell K, Mitra RD. 2019. Homotypic cooperativity and collective binding are determinants of bHLH specificity and function. Proc. Natl. Acad. Sci. U. S. A. 116:16143–16152.

Shubin N, Tabin C, Carroll S. 1997. Fossils, genes and the evolution of animal limbs. Nature 388:639–648.

Siddiq MA, Duveau F, Wittkopp PJ. 2024. Plasticity and environment-specific relationships between gene expression and fitness in Saccharomyces cerevisiae. Nat. Ecol. Evol.:1–11.

Stern DL, Orgogozo V. 2008. The loci of evolution: how predictable is genetic evolution? Evolution 62:2155–2177.

Tornow J, Zeng X, Gao W, Santangelo GM. 1993. GCR1, a transcriptional activator in Saccharomyces cerevisiae, complexes with RAP1 and can function without its DNA binding domain. EMBO J. 12:2431–2437.

Uemura H, Jigami Y. 1992. Role of GCR2 in transcriptional activation of yeast glycolytic genes. Mol. Cell. Biol. 12:3834–3842.

Umans BD, Battle A, Gilad Y. 2021. Where are the disease-associated eQTLs? Trends Genet. 37:109–124.

Vaishnav ED, de Boer CG, Molinet J, Yassour M, Fan L, Adiconis X, Thompson DA, Levin JZ, Cubillos FA, Regev A. 2022. The evolution, evolvability and engineering of gene regulatory DNA. Nature 603:455–463.

Vande Zande P, Hill MS, Wittkopp PJ. 2022. Pleiotropic effects of trans-regulatory mutations on fitness and gene expression. Science 377:105–109.

Vande Zande P, Siddiq MA, Hodgins-Davis A, Kim L, Wittkopp PJ. 2023. Active compensation for changes in TDH3 expression mediated by direct regulators of TDH3 in Saccharomyces cerevisiae. PLoS Genet. 19:e1011078.

Wittkopp PJ. 2023. Contributions of mutation and selection to regulatory variation: lessons from the Saccharomyces cerevisiae TDH3 gene. Philos. Trans. R. Soc. Lond. B Biol. Sci. 378:20220057.

Wray GA. 2007. The evolutionary significance of cis-regulatory mutations. Nat. Rev. Genet. 8:206–216.

Xie Z, Sokolov I, Osmala M, Yue X, Bower G, Pett JP, Chen Y, Wang K, Cavga AD, Popov A, et al. 2025. DNA-guided transcription factor interactions extend human gene regulatory code. Nature 641:1329–1338.

