## Supplementary File 1 for "*cis-* and *trans*-regulatory factors contributing to divergent activity of the *TDH3* promoter in *Saccharomyces* yeast"

**Supplementary File 1 for “*cis*- and *trans*-regulatory factors contributing to divergent activity of the *TDH3* promoter in *Saccharomyces* yeast” by Mohammad A. Siddiq, Hannah P. Kania, Nicholas J. Brown, and Patricia J. Wittkopp**

Sequences of the integrated reporter constructs at the HO locus. The sequences of the orthologous promoters are shown in different colors(S. cer: Blue; S. par: Green ; S. mik: pink; S. kud: brown). Venus YFP is highlighted in yellow. The cyc1 terminator sequence is shown in gray. The KanMX cassette is underlined.

>Scer_Ptdh3-YFP-KAN

CATCCAAAATATTAAATTTTACTTTTATTACATACAACTTTTTAAACTAATATACACATTGTTCGAGTTTATCATTATCAATACTGCCATTTCAAAGAATACGTAAATAATTAATAGTAGTGATTTTCCTAACTTTATTTAGTCAAAAAATTAGCCTTTTAATTCTGCTGTAACCCGTACATGCCCAAAATAGGGGGCGGGTTACACAGAATATATAACATCGTAGGTGTCTGGGTGAACAGTTTATTCCTGGCATCCACTAAATATAATGGAGCCCGCTTTTTAAGCTGGCATCCAGaaaaaaaaaGAATCCCAGCACCAAAATATTGTTTTCTTCACCAACCATCAGTTCATAGGTCCATTCTCTTAGCGCAACTACAGAGAACAGGGGCACAAACAGGCAAAAAACGGGCACAACCTCAATGGAGTGATGCAACCTGCCTGGAGTAAATGATGACACAAGGCAATTGACCCACGCATGTATCTATCTCATTTTCTTACACCTTCTATTACCTTCTGCTCTCTCTGATTTGGaaaaagctgaaaaaaaaGGTTGAAACCAGTTCCCTGAAATTATTCCCCTACTTGACTAATAAGTATATAAAGACGGTAGGTATTGATTGTAATTCTGTAAATCTATTTCTTAAACTTCTTAAATTCTACTTTTATAGTTAGTCttttttttagttttAAAACACCAAGAACTTAGTTTCGAATAAACACACATAAACAAACAAAATGTCTAAAGGTGAAGAATTATTCACTGGTGTTGTCCCAATTTTGGTTGAATTAGATGGTGATGTTAATGGTCACAAATTTTCTGTCTCCGGTGAAGGTGAAGGTGATGCTACTTACGGTAAATTGACCTTAAAATTAATTTGTACTACTGGTAAATTGCCAGTTCCATGGCCAACCTTAGTCACTACTTTAGGTTATGGTTTAATGTGTTTTGCTAGATACCCAGATCATATGAAACAACATGACTTTTTCAAGTCTGCCATGCCAGAAGGTTATGTTCAAGAAAGAACTATTTTTTTCAAAGATGACGGTAACTACAAGACCAGAGCTGAAGTCAAGTTTGAAGGTGATACCTTAGTTAATAGAATCGAATTAAAAGGTATTGATTTTAAAGAAGATGGTAACATTTTAGGTCACAAATTGGAATACAACTATAACTCTCACAATGTTTACATCACTGCTGACAAACAAAAGAATGGTATCAAAGCTAACTTCAAAATTAGACACAACATTGAAGATGGTGGTGTTCAATTAGCTGACCATTATCAACAAAATACTCCAATTGGTGATGGTCCAGTCTTGTTACCAGACAACCATTACTTATCCTATCAATCTGCCTTATCCAAAGATCCAAACGAAAAGAGAGACCACATGGTCTTGTTAGAATTTGTTACTGCTGCTGGTATTACCCATGGTATGGATGAATTGTACAAATAGTTCCTTTGTCGATATCATGTAATTAGTTATGTCACGCTTACATTCACGCCCTCCTCCCACATCCGCTCTAACCGAAAAGGAAGGAGTTAGACAACCTGAAGTCTAGGTCCCTATTTATTTTTTTTAATAGTTATGTTAGTATTAAGAACGTTATTTATATCTCAAATTTTTCTTTTTTTTCTGTACAAACGCGTGTACGCATGTAACATTATACTGAAAACCTTGCTTGAGAAGGTTTTGGGACGCTCGAAGGCTTTAATTTGCAAGCTTCGCGTTTACACTTGGCAAAGCAAGGTCCATGTTTTCTTGAAAGGTAAGAAGTGATCGATGAATTCGAGCTCGTTTTCGACACTGGATGGCGGCGTTAGTATCGAATCGACAGCAGTATAGCGACCAGCATTCACATACGATTGACGCATGATATTACTTTCTGCGCACTTAACTTCGCATCTGGGCAGATGATGTCGAGGCGAAAAAAAATATAAATCACGCTAACATTTGATTAAAATAGAACAACTACAATATAAAAAAACTATACAAATGACAAGTTCTTGAAAACAAGAATCTTTTTATTGTCAGTACTGATTAGAAAAACTCATCGAGCATCAAATGAAACTGCAATTTATTCATATCAGGATTATCAATACCATATTTTTGAAAAAGCCGTTTCTGTAATGAAGGAGAAAACTCACCGAGGCAGTTCCATAGGATGGCAAGATCCTGGTATCGGTCTGCGATTCCGACTCGTCCAACATCAATACAACCTATTAATTTCCCCTCGTCAAAAATAAGGTTATCAAGTGAGAAATCACCATGAGTGACGACTGAATCCGGTGAGAATGGCAAAAGCTTATGCATTTCTTTCCAGACTTGTTCAACAGGCCAGCCATTACGCTCGTCATCAAAATCACTCGCATCAACCAAACCGTTATTCATTCGTGATTGCGCCTGAGCGAGACGAAATACGCGATCGCTGTTAAAAGGACAATTACAAACAGGAATCGAATGCAACCGGCGCAGGAACACTGCCAGCGCATCAACAATATTTTCACCTGAATCAGGATATTCTTCTAATACCTGGAATGCTGTTTTGCCGGGGATCGCAGTGGTGAGTAACCATGCATCATCAGGAGTACGGATAAAATGCTTGATGGTCGGAAGAGGCATAAATTCCGTCAGCCAGTTTAGTCTGACCATCTCATCTGTAACATCATTGGCAACGCTACCTTTGCCATGTTTCAGAAACAACTCTGGCGCATCGGGCTTCCCATACAATCGATAGATTGTCGCACCTGATTGCCCGACATTATCGCGAGCCCATTTATACCCATATAAATCAGCATCCATGTTGGAATTTAATCGCGGCCTCGAAACGTGAGTCTTTTCCTTACCCATGGTTGTTTATGTTCGGATGTGATGTGAGAACTGTATCCTAGCAAGATTTTAAAAGGAAGTATATGAAAGAAGAACCTCAGTGGCAAATCCTAACCTTTTATATTTCTCTACAGGGGCGCGGCGTGGGGACAATTCAACGCGTCTGTGAGGGGAGCGTTTCCCTGCTCGCAGGTCTGCAGCGAGGAGCCGTAATTTTTGCTTCGCGCCGTGCGGCCATCAAAATGTATGGATGCAAATGATTATACATGGGGATGTATGGGCTAAATGTACGGGCGACAGTCACATCATGCCCCTGAGCTGCGCACGTCAAGACTGTCAAGGAGGGTATTCTGGGCCTCCATGTCGCTGGCCGGGTGACCCGGCGGGGACGAGGCAAGC*T*AAACAGATCTGGCGCGCCTTAATTAACCCGGGGATCCGTCGAC

>Spar_Ptdh3-YFP-KAN

CATCCAAAATATTAAATTTTACTTTTATTACATACAACTTTTTAAACTAATATACACATTGTTCAAATTTATCGTTGTCATTACCACCATTTTAAAGAATACGTAAATAGTTAATAGTAATGATTTTCCTAACTTTATTAAGTCAAAAAATTTACGTTTTATTTCTGCTGCCATCCGTAAATGCCAGGATTTGAGCGGGTTACACAATATATCTCATATTTTCGGTGTCTGGGTCATTACTTTACTCTTGGCATCCACTAAATATATTGGATCCTGCTTTTTAAACTGGCTTCCAGAAAAAAAAAGAGAATCCCAGCACCAAAAGATAGTTTTCCTCACTAACCGTCAGATCATAGGTCCATTCTCTTAGCGCAACTACACAGAACAGGGGCACAAACAGGCAAAAAACGGGCACAACCTCAATGGAGTGATGCAAACTGCCTGGAGTAAAAGATGACACAAGGCGATTGACCTACGCATGTATCTATCTCATTTTCTTACACCTTCTATTTCATTCTAACTCTTTGATTTGGAAAACACCTAAGAAAAAAAAGGTTGAAATCAGTTCCCTGAAATTGTCCCCCTACTTGACTAATAAATATATAAAGACGGTAGGTATTGACTGTAATTCGTAAATCTATACTTCTTAAACTTCTTCAAATTTACTTTTTTGGATAGTCTTATTTTTGGTTTCAATACCCCAAGAACTTAGTTTCAAATAAATACACATACAAACAAAATGTCTAAAGGTGAAGAATTATTCACTGGTGTTGTCCCAATTTTGGTTGAATTAGATGGTGATGTTAATGGTCACAAATTTTCTGTCTCCGGTGAAGGTGAAGGTGATGCTACTTACGGTAAATTGACCTTAAAATTAATTTGTACTACTGGTAAATTGCCAGTTCCATGGCCAACCTTAGTCACTACTTTAGGTTATGGTTTAATGTGTTTTGCTAGATACCCAGATCATATGAAACAACATGACTTTTTCAAGTCTGCCATGCCAGAAGGTTATGTTCAAGAAAGAACTATTTTTTTCAAAGATGACGGTAACTACAAGACCAGAGCTGAAGTCAAGTTTGAAGGTGATACCTTAGTTAATAGAATCGAATTAAAAGGTATTGATTTTAAAGAAGATGGTAACATTTTAGGTCACAAATTGGAATACAACTATAACTCTCACAATGTTTACATCACTGCTGACAAACAAAAGAATGGTATCAAAGCTAACTTCAAAATTAGACACAACATTGAAGATGGTGGTGTTCAATTAGCTGACCATTATCAACAAAATACTCCAATTGGTGATGGTCCAGTCTTGTTACCAGACAACCATTACTTATCCTATCAATCTGCCTTATCCAAAGATCCAAACGAAAAGAGAGACCACATGGTCTTGTTAGAATTTGTTACTGCTGCTGGTATTACCCATGGTATGGATGAATTGTACAAATAGTTCCTTTGTCGATATCATGTAATTAGTTATGTCACGCTTACATTCACGCCCTCCTCCCACATCCGCTCTAACCGAAAAGGAAGGAGTTAGACAACCTGAAGTCTAGGTCCCTATTTATTTTTTTTAATAGTTATGTTAGTATTAAGAACGTTATTTATATCTCAAATTTTTCTTTTTTTTCTGTACAAACGCGTGTACGCATGTAACATTATACTGAAAACCTTGCTTGAGAAGGTTTTGGGACGCTCGAAGGCTTTAATTTGCAAGCTTCGCGTTTACACTTGGCAAAGCAAGGTCCATGTTTTCTTGAAAGGTAAGAAGTGATCGATGAATTCGAGCTCGTTTTCGACACTGGATGGCGGCGTTAGTATCGAATCGACAGCAGTATAGCGACCAGCATTCACATACGATTGACGCATGATATTACTTTCTGCGCACTTAACTTCGCATCTGGGCAGATGATGTCGAGGCGAAAAAAAATATAAATCACGCTAACATTTGATTAAAATAGAACAACTACAATATAAAAAAACTATACAAATGACAAGTTCTTGAAAACAAGAATCTTTTTATTGTCAGTACTGATTAGAAAAACTCATCGAGCATCAAATGAAACTGCAATTTATTCATATCAGGATTATCAATACCATATTTTTGAAAAAGCCGTTTCTGTAATGAAGGAGAAAACTCACCGAGGCAGTTCCATAGGATGGCAAGATCCTGGTATCGGTCTGCGATTCCGACTCGTCCAACATCAATACAACCTATTAATTTCCCCTCGTCAAAAATAAGGTTATCAAGTGAGAAATCACCATGAGTGACGACTGAATCCGGTGAGAATGGCAAAAGCTTATGCATTTCTTTCCAGACTTGTTCAACAGGCCAGCCATTACGCTCGTCATCAAAATCACTCGCATCAACCAAACCGTTATTCATTCGTGATTGCGCCTGAGCGAGACGAAATACGCGATCGCTGTTAAAAGGACAATTACAAACAGGAATCGAATGCAACCGGCGCAGGAACACTGCCAGCGCATCAACAATATTTTCACCTGAATCAGGATATTCTTCTAATACCTGGAATGCTGTTTTGCCGGGGATCGCAGTGGTGAGTAACCATGCATCATCAGGAGTACGGATAAAATGCTTGATGGTCGGAAGAGGCATAAATTCCGTCAGCCAGTTTAGTCTGACCATCTCATCTGTAACATCATTGGCAACGCTACCTTTGCCATGTTTCAGAAACAACTCTGGCGCATCGGGCTTCCCATACAATCGATAGATTGTCGCACCTGATTGCCCGACATTATCGCGAGCCCATTTATACCCATATAAATCAGCATCCATGTTGGAATTTAATCGCGGCCTCGAAACGTGAGTCTTTTCCTTACCCATGGTTGTTTATGTTCGGATGTGATGTGAGAACTGTATCCTAGCAAGATTTTAAAAGGAAGTATATGAAAGAAGAACCTCAGTGGCAAATCCTAACCTTTTATATTTCTCTACAGGGGCGCGGCGTGGGGACAATTCAACGCGTCTGTGAGGGGAGCGTTTCCCTGCTCGCAGGTCTGCAGCGAGGAGCCGTAATTTTTGCTTCGCGCCGTGCGGCCATCAAAATGTATGGATGCAAATGATTATACATGGGGATGTATGGGCTAAATGTACGGGCGACAGTCACATCATGCCCCTGAGCTGCGCACGTCAAGACTGTCAAGGAGGGTATTCTGGGCCTCCATGTCGCTGGCCGGGTGACCCGGCGGGGACGAGGCAAGC*T*AAACAGATCTGGCGCGCCTTAATTAACCCGGGGATCCGTCGAC

>Smik_Ptdh3-YFP-KAN

CATCCAAAATATTAAATTTTACTTTTATTACATACAACTTTTTAAACTAATATACACATTCTTCTAATTTATCGTTGTCATTACAACCATTTGAAAAAATACGTAAATAGCCAATAGTAATGATTTTCCTAACCTTATTAAGCTGTAACACAAGTACTGCTATTTTTCCATTATTCCTACCCATCCAATCATACGGCGGGTTACACATCCACTCGCCATTTTAGATGTCTGGGTGATTGATTATATCACCGGCATCCACTAATCAGATGGATCCCGCTTTTTATGTTGGCTTCCAGAAAAAAAAAAAAGAATCCCAGCACCAAAAGGTACCTTTCTTCAACAACCATCAGATCATAGGTCCATTCTCTTACCGCAACCACACAGAACAGGGGCACAAACAGGCAAAAAGCGGACACAACCTCAATGGAGTGATGCAAACTGTCTGGAGCAAAAGAAGACACAAGGCATTTGAACTACGCATGTATCTAATCTTCTTACACCTTCTACTACCTTCTAACTCTTTGGGTTGGAAAAAAATGAAAAAAATAAATTGGAGTTAGTTCCCTGAAATTATCCCCCTACTTGACTAGTAAGTATATAAAGACGGTAAGTATTGATCATATTTCTGTAAATCTCTATTCATTAAGCTTCTTTACTTCGTATTAATTTTATTTAATCTTTTTAGTTAAATCCCCAAGAACTTAGTTTCGAATAAACACACACAAACAAACAAAATGTCTAAAGGTGAAGAATTATTCACTGGTGTTGTCCCAATTTTGGTTGAATTAGATGGTGATGTTAATGGTCACAAATTTTCTGTCTCCGGTGAAGGTGAAGGTGATGCTACTTACGGTAAATTGACCTTAAAATTAATTTGTACTACTGGTAAATTGCCAGTTCCATGGCCAACCTTAGTCACTACTTTAGGTTATGGTTTAATGTGTTTTGCTAGATACCCAGATCATATGAAACAACATGACTTTTTCAAGTCTGCCATGCCAGAAGGTTATGTTCAAGAAAGAACTATTTTTTTCAAAGATGACGGTAACTACAAGACCAGAGCTGAAGTCAAGTTTGAAGGTGATACCTTAGTTAATAGAATCGAATTAAAAGGTATTGATTTTAAAGAAGATGGTAACATTTTAGGTCACAAATTGGAATACAACTATAACTCTCACAATGTTTACATCACTGCTGACAAACAAAAGAATGGTATCAAAGCTAACTTCAAAATTAGACACAACATTGAAGATGGTGGTGTTCAATTAGCTGACCATTATCAACAAAATACTCCAATTGGTGATGGTCCAGTCTTGTTACCAGACAACCATTACTTATCCTATCAATCTGCCTTATCCAAAGATCCAAACGAAAAGAGAGACCACATGGTCTTGTTAGAATTTGTTACTGCTGCTGGTATTACCCATGGTATGGATGAATTGTACAAATAGTTCCTTTGTCGATATCATGTAATTAGTTATGTCACGCTTACATTCACGCCCTCCTCCCACATCCGCTCTAACCGAAAAGGAAGGAGTTAGACAACCTGAAGTCTAGGTCCCTATTTATTTTTTTTAATAGTTATGTTAGTATTAAGAACGTTATTTATATCTCAAATTTTTCTTTTTTTTCTGTACAAACGCGTGTACGCATGTAACATTATACTGAAAACCTTGCTTGAGAAGGTTTTGGGACGCTCGAAGGCTTTAATTTGCAAGCTTCGCGTTTACACTTGGCAAAGCAAGGTCCATGTTTTCTTGAAAGGTAAGAAGTGATCGATGAATTCGAGCTCGTTTTCGACACTGGATGGCGGCGTTAGTATCGAATCGACAGCAGTATAGCGACCAGCATTCACATACGATTGACGCATGATATTACTTTCTGCGCACTTAACTTCGCATCTGGGCAGATGATGTCGAGGCGAAAAAAAATATAAATCACGCTAACATTTGATTAAAATAGAACAACTACAATATAAAAAAACTATACAAATGACAAGTTCTTGAAAACAAGAATCTTTTTATTGTCAGTACTGATTAGAAAAACTCATCGAGCATCAAATGAAACTGCAATTTATTCATATCAGGATTATCAATACCATATTTTTGAAAAAGCCGTTTCTGTAATGAAGGAGAAAACTCACCGAGGCAGTTCCATAGGATGGCAAGATCCTGGTATCGGTCTGCGATTCCGACTCGTCCAACATCAATACAACCTATTAATTTCCCCTCGTCAAAAATAAGGTTATCAAGTGAGAAATCACCATGAGTGACGACTGAATCCGGTGAGAATGGCAAAAGCTTATGCATTTCTTTCCAGACTTGTTCAACAGGCCAGCCATTACGCTCGTCATCAAAATCACTCGCATCAACCAAACCGTTATTCATTCGTGATTGCGCCTGAGCGAGACGAAATACGCGATCGCTGTTAAAAGGACAATTACAAACAGGAATCGAATGCAACCGGCGCAGGAACACTGCCAGCGCATCAACAATATTTTCACCTGAATCAGGATATTCTTCTAATACCTGGAATGCTGTTTTGCCGGGGATCGCAGTGGTGAGTAACCATGCATCATCAGGAGTACGGATAAAATGCTTGATGGTCGGAAGAGGCATAAATTCCGTCAGCCAGTTTAGTCTGACCATCTCATCTGTAACATCATTGGCAACGCTACCTTTGCCATGTTTCAGAAACAACTCTGGCGCATCGGGCTTCCCATACAATCGATAGATTGTCGCACCTGATTGCCCGACATTATCGCGAGCCCATTTATACCCATATAAATCAGCATCCATGTTGGAATTTAATCGCGGCCTCGAAACGTGAGTCTTTTCCTTACCCATGGTTGTTTATGTTCGGATGTGATGTGAGAACTGTATCCTAGCAAGATTTTAAAAGGAAGTATATGAAAGAAGAACCTCAGTGGCAAATCCTAACCTTTTATATTTCTCTACAGGGGCGCGGCGTGGGGACAATTCAACGCGTCTGTGAGGGGAGCGTTTCCCTGCTCGCAGGTCTGCAGCGAGGAGCCGTAATTTTTGCTTCGCGCCGTGCGGCCATCAAAATGTATGGATGCAAATGATTATACATGGGGATGTATGGGCTAAATGTACGGGCGACAGTCACATCATGCCCCTGAGCTGCGCACGTCAAGACTGTCAAGGAGGGTATTCTGGGCCTCCATGTCGCTGGCCGGGTGACCCGGCGGGGACGAGGCAAGC*T*AAACAGATCTGGCGCGCCTTAATTAACCCGGGGATCCGTCGAC

>Skud_Ptdh3-YFP-KAN

CATCCAAAATATTAAATTTTACTTTTATTACATACAACTTTTTAAACTAATATACACATTCCACGTATATATCGTTATCGTCGCTACCTCCTTAAAAAAATACGTAAATACTCGAAATAGTGATTTTCCTAACCTTATTAAATCAATTCATCGGCCCTTTTAGCGGCTACCCGCGCCATCTAGATGATAGGGCGGGTGACACTATGGTAAATCCCATAATTAGGTGTCTGGGTGAGTGGTTCTGATGCCGGCATCCACTAAATATATTGGATCCCATTTTTTACGCGGGCTTCCAGAAAAAAAGAGAATCCCAGCACCAAAAGGTGGTTCTCTTCACCAACCATCAGATCATAGGTCCACAACCACACATAACAGGGGCACAAAAAGGCAAAAAACGGACATAACCTCAATGGAGTGATGCAAATTGACTGGAGCAAAAGCTGACACAAGGCATTGATTGACCTACGCATGTATCTGTATTCTTTTCTTACACCTTCTATTACCTTCTAACTCTTTGGGTTGGAAAAAACTGAAAAAAAAGGTTGGGACCTGGTTCCCCCAAGTTGTCCCCCTACTTGGTTATTAAATATATAAAGACAGCAAGTGTTGATTATAATCTTGTAAATCTATAGTTCTTAATCTATACTTCTATTTATATTTTAAATTAGTCTTTTTATTTCCAAGTCCCCAAGAACTTAGTTTCGAATAAACACACACAAATAAACACATGTCTAAAGGTGAAGAATTATTCACTGGTGTTGTCCCAATTTTGGTTGAATTAGATGGTGATGTTAATGGTCACAAATTTTCTGTCTCCGGTGAAGGTGAAGGTGATGCTACTTACGGTAAATTGACCTTAAAATTAATTTGTACTACTGGTAAATTGCCAGTTCCATGGCCAACCTTAGTCACTACTTTAGGTTATGGTTTAATGTGTTTTGCTAGATACCCAGATCATATGAAACAACATGACTTTTTCAAGTCTGCCATGCCAGAAGGTTATGTTCAAGAAAGAACTATTTTTTTCAAAGATGACGGTAACTACAAGACCAGAGCTGAAGTCAAGTTTGAAGGTGATACCTTAGTTAATAGAATCGAATTAAAAGGTATTGATTTTAAAGAAGATGGTAACATTTTAGGTCACAAATTGGAATACAACTATAACTCTCACAATGTTTACATCACTGCTGACAAACAAAAGAATGGTATCAAAGCTAACTTCAAAATTAGACACAACATTGAAGATGGTGGTGTTCAATTAGCTGACCATTATCAACAAAATACTCCAATTGGTGATGGTCCAGTCTTGTTACCAGACAACCATTACTTATCCTATCAATCTGCCTTATCCAAAGATCCAAACGAAAAGAGAGACCACATGGTCTTGTTAGAATTTGTTACTGCTGCTGGTATTACCCATGGTATGGATGAATTGTACAAATAGTTCCTTTGTCGATATCATGTAATTAGTTATGTCACGCTTACATTCACGCCCTCCTCCCACATCCGCTCTAACCGAAAAGGAAGGAGTTAGACAACCTGAAGTCTAGGTCCCTATTTATTTTTTTTAATAGTTATGTTAGTATTAAGAACGTTATTTATATCTCAAATTTTTCTTTTTTTTCTGTACAAACGCGTGTACGCATGTAACATTATACTGAAAACCTTGCTTGAGAAGGTTTTGGGACGCTCGAAGGCTTTAATTTGCAAGCTTCGCGTTTACACTTGGCAAAGCAAGGTCCATGTTTTCTTGAAAGGTAAGAAGTGATCGATGAATTCGAGCTCGTTTTCGACACTGGATGGCGGCGTTAGTATCGAATCGACAGCAGTATAGCGACCAGCATTCACATACGATTGACGCATGATATTACTTTCTGCGCACTTAACTTCGCATCTGGGCAGATGATGTCGAGGCGAAAAAAAATATAAATCACGCTAACATTTGATTAAAATAGAACAACTACAATATAAAAAAACTATACAAATGACAAGTTCTTGAAAACAAGAATCTTTTTATTGTCAGTACTGATTAGAAAAACTCATCGAGCATCAAATGAAACTGCAATTTATTCATATCAGGATTATCAATACCATATTTTTGAAAAAGCCGTTTCTGTAATGAAGGAGAAAACTCACCGAGGCAGTTCCATAGGATGGCAAGATCCTGGTATCGGTCTGCGATTCCGACTCGTCCAACATCAATACAACCTATTAATTTCCCCTCGTCAAAAATAAGGTTATCAAGTGAGAAATCACCATGAGTGACGACTGAATCCGGTGAGAATGGCAAAAGCTTATGCATTTCTTTCCAGACTTGTTCAACAGGCCAGCCATTACGCTCGTCATCAAAATCACTCGCATCAACCAAACCGTTATTCATTCGTGATTGCGCCTGAGCGAGACGAAATACGCGATCGCTGTTAAAAGGACAATTACAAACAGGAATCGAATGCAACCGGCGCAGGAACACTGCCAGCGCATCAACAATATTTTCACCTGAATCAGGATATTCTTCTAATACCTGGAATGCTGTTTTGCCGGGGATCGCAGTGGTGAGTAACCATGCATCATCAGGAGTACGGATAAAATGCTTGATGGTCGGAAGAGGCATAAATTCCGTCAGCCAGTTTAGTCTGACCATCTCATCTGTAACATCATTGGCAACGCTACCTTTGCCATGTTTCAGAAACAACTCTGGCGCATCGGGCTTCCCATACAATCGATAGATTGTCGCACCTGATTGCCCGACATTATCGCGAGCCCATTTATACCCATATAAATCAGCATCCATGTTGGAATTTAATCGCGGCCTCGAAACGTGAGTCTTTTCCTTACCCATGGTTGTTTATGTTCGGATGTGATGTGAGAACTGTATCCTAGCAAGATTTTAAAAGGAAGTATATGAAAGAAGAACCTCAGTGGCAAATCCTAACCTTTTATATTTCTCTACAGGGGCGCGGCGTGGGGACAATTCAACGCGTCTGTGAGGGGAGCGTTTCCCTGCTCGCAGGTCTGCAGCGAGGAGCCGTAATTTTTGCTTCGCGCCGTGCGGCCATCAAAATGTATGGATGCAAATGATTATACATGGGGATGTATGGGCTAAATGTACGGGCGACAGTCACATCATGCCCCTGAGCTGCGCACGTCAAGACTGTCAAGGAGGGTATTCTGGGCCTCCATGTCGCTGGCCGGGTGACCCGGCGGGGACGAGGCAAGC*T*AAACAGATCTGGCGCGCCTTAATTAACCCGGGGATCCGTCGAC
