## Supplementary File 2 for "*cis-* and *trans*-regulatory factors contributing to divergent activity of the *TDH3* promoter in *Saccharomyces* yeast"

**Supplementary File 2 for “*cis*- and *trans*-regulatory factors contributing to divergent activity of the *TDH3* promoter in *Saccharomyces* yeast” by Mohammad A. Siddiq, Hannah P. Kania, Nicholas J. Brown, and Patricia J. Wittkopp**

Sequences of the *TDH3* and *Tye7* loci in reference and deletion strains. Open reading frames corresponding to TDH3p and Tye7p are shown in bold. In the TDH3 deletion strain, the deleted bases are struck through. In the Tye7 deletion strain, the entire sequence of the Tye7 open reading frame was removed and replaced with the mScarlet-I fluorophore (highlighted in red). A table of strain sequence numbers is provided at the end of the file.

>Scer_Ref_TDH3

TTTCAGTTCGAGTTTATCATTATCAATACTGCCATTTCAAAGAATACGTAAATAATTAATAGTAGTGATTTTCCTAACTTTATTTAGTCAAAAAATTAGCCTTTTAATTCTGCTGTAACCCGTACATGCCCAAAATAGGGGGCGGGTTACACAGAATATATAACATCGTAGGTGTCTGGGTGAACAGTTTATTCCTGGCATCCACTAAATATAATGGAGCCCGCTTTTTAAGCTGGCATCCAGAAAAAAAAAGAATCCCAGCACCAAAATATTGTTTTCTTCACCAACCATCAGTTCATAGGTCCATTCTCTTAGCGCAACTACAGAGAACAGGGGCACAAACAGGCAAAAAACGGGCACAACCTCAATGGAGTGATGCAACCTGCCTGGAGTAAATGATGACACAAGGCAATTGACCCACGCATGTATCTATCTCATTTTCTTACACCTTCTATTACCTTCTGCTCTCTCTGATTTGGAAAAAGCTGAAAAAAAAGGTTGAAACCAGTTCCCTGAAATTATTCCCCTACTTGACTAATAAGTATATAAAGACGGTAGGTATTGATTGTAATTCTGTAAATCTATTTCTTAAACTTCTTAAATTCTACTTTTATAGTTAGTCTTTTTTTTAGTTTTAAAACACCAAGAACTTAGTTTCGAATAAACACACATAAACAAACAAA**ATGGTTAGAGTTGCTATTAACGGTTTCGGTAGAATCGGTAGATTGGTCATGAGAATTGCTTTGTCTAGACCAAACGTCGAAGTTGTTGCTTTGAACGACCCATTCATCACCAACGACTACGCTGCTTACATGTTCAAGTACGACTCCACTCACGGTAGATACGCTGGTGAAGTTTCCCACGATGACAAGCACATCATTGTCGATGGTAAGAAGATTGCTACTTACCAAGAAAGAGACCCAGCTAACTTGCCATGGGGTTCTTCCAACGTTGACATCGCCATTGACTCCACTGGTGTTTTCAAGGAATTAGACACTGCTCAAAAGCACATTGACGCTGGTGCCAAGAAGGTTGTTATCACTGCTCCATCTTCCACCGCCCCAATGTTCGTCATGGGTGTTAACGAAGAAAAATACACTTCTGACTTGAAGATTGTTTCCAACGCTTCTTGTACCACCAACTGTTTGGCTCCATTGGCCAAGGTTATCAACGATGCTTTCGGTATTGAAGAAGGTTTGATGACCACTGTCCACTCTTTGACTGCTACTCAAAAGACTGTTGACGGTCCATCCCACAAGGACTGGAGAGGTGGTAGAACCGCTTCCGGTAACATCATCCCATCCTCCACCGGTGCTGCTAAGGCTGTCGGTAAGGTCTTGCCAGAATTGCAAGGTAAGTTGACCGGTATGGCTTTCAGAGTCCCAACCGTCGATGTCTCCGTTGTTGACTTGACTGTCAAGTTGAACAAGGAAACCACCTACGATGAAATCAAGAAGGTTGTTAAGGCTGCCGCTGAAGGTAAGTTGAAGGGTGTTTTGGGTTACACCGAAGACGCTGTTGTCTCCTCTGACTTCTTGGGTGACTCTCACTCTTCCATCTTCGATGCTTCCGCTGGTATCCAATTGTCTCCAAAGTTCGTCAAGTTGGTCTCCTGGTACGACAACGAATACGGTTACTCTACCAGAGTTGTCGACTTGGTTGAACACGTTGCCAAGGCTTAA**GTGAATTTACTTTAAATCTTGCATTTAAATAAATTTTCTTTTTATAGCTTTATGACTTAGTTTCAATTTATATACTATTTTAATGACATTTTCGATTCATTGATTGAAAGCTTTGTGTTTTTTCTTGATGCGCTATTGCATTGTTCTTGTCTTTTTCGCCACATGTAATATCTGTAGTAGATACCTGATACATT

>Scer_del_TDH3

TTTCAGTTCGAGTTTATCATTATCAATACTGCCATTTCAAAGAATACGTAAATAATTAATAGTAGTGATTTTCCTAACTTTATTTAGTCAAAAAATTAGCCTTTTAATTCTGCTGTAACCCGTACATGCCCAAAATAGGGGGCGGGTTACACAGAATATATAACATCGTAGGTGTCTGGGTGAACAGTTTATTCCTGGCATCCACTAAATATAATGGAGCCCGCTTTTTAAGCTGGCATCCAGAAAAAAAAAGAATCCCAGCACCAAAATATTGTTTTCTTCACCAACCATCAGTTCATAGGTCCATTCTCTTAGCGCAACTACAGAGAACAGGGGCACAAACAGGCAAAAAACGGGCACAACCTCAATGGAGTGATGCAACCTGCCTGGAGTAAATGATGACACAAGGCAATTGACCCACGCATGTATCTATCTCATTTTCTTACACCTTCTATTACCTTCTGCTCTCTCTGATTTGGAAAAAGCTGAAAAAAAAGGTTGAAACCAGTTCCCTGAAATTATTCCCCTACTTGACTAATAAGTATATAAAGACGGTAGGTATTGATTGTAATTCTGTAAATCTATTTCTTAAACTTCTTAAATTCTACTTTTATAGTTAGTC~~TTTTTTTTAGTTTTAAAACACCAAGAACTTAGTTTCGAATAAACACACATAAACAAACAAA~~**~~ATGGTTAGAGTTGCTATTAACGGTTTCGGTAGAATCGGTAGATTGGTCATGAGAATTGCTTTGTCTAGACCAAACGTCGAAGTTGTTGCTTTGAACGACCCATTCATCACCAACGACTACGCTGCTTACATGTTCAAGTACGACTCCACTCACGGTAGATACGCTGGTGAAGTTTCCCACGATGACAAGCACATCATTGTCGATGGTAAGAAGATTGCTACTTACCAAGAAAGAGACCCAGCTAACTTGCCATGGGGTTCTTCCAACGTTGACATCGCCATTGACTCCACTGGTGTTTTCAAGGAATTAGACACTGCTCAAAAGCACATTGACGCTGGTGCCAAGAAGGTTGTTATCACTGCTCCATCTTCCACCGCCCCAATGTTCGTCATGGGTGTTAACGAAGAAAAATACACTTCTGACTTGAAGATTGTTTCCAACGCTTCTTGTACCACCAACTGTTTGGCTCCATTGGCCAAGGTTATCAACGATGCTTTCGGTATTGAAGAAGGTTTGATGACCACTGTCCACTCTTTGACTGCTACTCAAAAGACTGTTGACGGTCCATCCCACAAGGACTGGAGAGGTGGTAGAACCGCTTCCGGTAACATCATCCCATCCTCCACCGGTGCTGCTAAGGCTGTCGGTAAGGTCTTGCCAGAATTGCAAGGTAAGTTGACCGGTATGGCTTTCAGAGTCCCAACCGTCGATGTCTCCGTTGTTGACTTGACTGTCAAGTTGAACAAGGAAACCACCTACGATGAAATCAAGAAGGTTGTTAAGGCTGCCGCTGAAGGTAAGTTGAAGGGTGTTTTGGGTTACACCGAAGACGCTGTTGTCTCCTCTGACTTCTTGGGTGACTCTCACTCTTCCATCTTCGATGCTTCCGCTGGTATCCAATTGTCTCCAAAGTTCGTCAAGTTGGTCTCCTGGTACGACAACGAATACGGTTACTCTACCAGAGTTGTCGACTTGGTTGAACACGTTGCCAAGGCTTAA~~**~~GTGAATTTACTTTAAATCTTGCATTTAAATAAATTTTCTTTTTATAGCTTTATGACTTAGTTTCAATTTATATACTATTTTAATGACATTTTCGATTCATTGATTGAAAGCTTTGTGTTTTTT~~CTTGATGCGCTATTGCATTGTTCTTGTCTTTTTCGCCACATGTAATATCTGTAGTAGATACCTGATACATT

>Scer_Ref_Tye7

TCTGCCCTGCGCACGTTCACAGTTCGCCAAATAACAATTCTCATTTTCATTTCTTTTTCACAGTGTAGATGATACCGTGTCTGCGACATTTCCATTCTTAGTAGAAATGCAAAACGCAACGTTTGCTATTAATTTTTTTCACCTCTGTTTCCTTTTTTTTTCTCACCTCAACAATCAAATGTACTTATATAAACTTCTTTTGTTATCTCCAAAATTTAAACTTATTTACATCCTTTTTCACTTCATATATCAATTCATGGGAAAGTAAAGCTATCTTAATTACATCCTTATTTTATATTTCCCAACCATCAACCTTTATTCTTTACATATTTCTCTCTTTTCTCATTTTCTTTCTTATTAAGCGTAAAGCACCACAAGCAGTACCGATCTAAAGTCTCTCTACTTTACTATTATTTTTTTTTTTTGTTCACTTCATAAACTTTTAGTGCAAAAATAAAAATAAAGACAAATAACACACTATCAAATCTCTTCAAGTTTAACAATT**ATGAACTCTATTTTAGACAGAAATGTTAGATCTAGCGAAACTACTTTAATTAAACCTGAATCTGAATTTGATAATTGGTTGTCGGATGAAAATGACGGAGCTAGTCATATCAACGTCAACAAGGACTCCTCGTCAGTTCTTTCTGCATCTTCTTCCACATGGTTCGAACCATTGGAAAACATTATCTCCTCTGCATCCAGCTCCTCGATAGGCTCTCCAATCGAAGACCAGTTTATATCTTCCAACAACGAGGAATCTGCTCTTTTTCCAACAGATCAGTTTTTCAGTAATCCTTCCTCATACTCGCATTCTCCCGAGGTTAGCAGCTCGATAAAAAGAGAAGAGGATGACAATGCCCTTTCGTTGGCAGATTTTGAACCGGCTTCTTTGCAATTAATGCCTAACATGATAAATACTGATAATAATGACGATAGTACCCCACTTAAGAATGAAATCGAGCTAAACGACTCGTTTATAAAAACAAATCTAGATGCTAAGGAAACGAAAAAGAGGGCTCCAAGAAAAAGACTGACCCCCTTCCAAAAGCAAGCTCACAACAAGATTGAAAAACGCTACAGAATAAACATCAACACAAAGATTGCAAGACTGCAGCAGATTATCCCATGGGTAGCAAGTGAACAAACAGCTTTCGAAGTAGGTGATTCTGTAAAAAAACAGGACGAAGACGGCGCAGAAACTGCCGCTACTACTCCTCTTCCATCTGCCGCTGCTACAAGCACGAAGCTAAATAAAAGCATGATCCTAGAAAAAGCTGTTGACTATATTCTATATCTACAAAATAACGAACGACTATACGAAATGGAAGTTCAAAGGTTGAAAAGTGAAATCGACACTTTGAAACAAGACCAAAAATAA**AAGCACGCTTGCTTAAACACCAATATTATAAAATTACGATAGAAATTCCTTTCTCCTCTTTTGTTTATGCGACATTTTTTTCTTTCTTAGCGAAGTACTTTAAAAAACAAGCCCAAAAAATAACAAGGCCATAACATAGCA

>Scer_del_Tye7

TCTGCCCTGCGCACGTTCACAGTTCGCCAAATAACAATTCTCATTTTCATTTCTTTTTCACAGTGTAGATGATACCGTGTCTGCGACATTTCCATTCTTAGTAGAAATGCAAAACGCAACGTTTGCTATTAATTTTTTTCACCTCTGTTTCCTTTTTTTTTCTCACCTCAACAATCAAATGTACTTATATAAACTTCTTTTGTTATCTCCAAAATTTAAACTTATTTACATCCTTTTTCACTTCATATATCAATTCATGGGAAAGTAAAGCTATCTTAATTACATCCTTATTTTATATTTCCCAACCATCAACCTTTATTCTTTACATATTTCTCTCTTTTCTCATTTTCTTTCTTATTAAGCGTAAAGCACCACAAGCAGTACCGATCTAAAGTCTCTCTACTTTACTATTATTTTTTTTTTTTGTTCACTTCATAAACTTTTAGTGCAAAAATAAAAATAAAGACAAATAACACACTATCAAATCTCTTCAAGTTTAACAATT**ATGGTTAGTAAAGGTGAAGCTGTTATAAAAGAATTCATGAGGTTTAAAGTTCATATGGAAGGTTCAATGAATGGTCATGAATTTGAAATTGAAGGTGAAGGTGAAGGTAGACCATATGAAGGTACACAAACTGCTAAATTGAAAGTTACTAAAGGTGGTCCATTACCATTTTCTTGGGATATTTTGTCTCCACAATTCATGTATGGTTCTAGAGCTTTTATTAAGCATCCAGCTGATATTCCAGATTATTATAAACAATCTTTTCCTGAAGGTTTTAAATGGGAAAGAGTTATGAATTTCGAAGATGGTGGTGCTGTTACTGTTACTCAAGATACTTCTTTGGAAGATGGTACTTTAATCTATAAAGTTAAATTGAGAGGTACTAATTTTCCACCAGATGGTCCAGTTATGCAAAAGAAAACTATGGGTTGGGAAGCATCTACTGAAAGATTGTATCCAGAAGATGGTGTTTTGAAAGGAGACATTAAGATGGCTTTGAGATTGAAAGATGGTGGTCGGTACCTGGCCGACTTCAAGACCACCTATAAAGCTAAAAAACCAGTTCAAATGCCAGGTGCATATAATGTTGATAGAAAGTTAGATATAACGTCGCATAACGAGGACTATACTGTAGTTGAACAATATGAACGTAGTGAAGGTAGACATAGTACCGGAGGAATGGATGAATTGTATAA**AAGCACGCTTGCTTAAACACCAATATTATAAAATTACGATAGAAATTCCTTTCTCCTCTTTTGTTTATGCGACATTTTTTTCTTTCTTAGCGAAGTACTTTAAAAAACAAGCCCAAAAAATAACAAGGCCATAACATAGCA

| Strain # | HO Reporter Construct | TDH3 genotype | Tye7 genotype |
| --- | --- | --- | --- |
| Y3326&Y3986 | None | Reference | Reference |
| Y3324&Y3872 | ScerPtdh3-YFP-cyc1 | Reference | Reference |
| Y3860 | SmikPtdh3-YFP-cyc1 | Reference | Reference |
| Y3861 | SkudPtdh3-YFP-cyc1 | Reference | Reference |
| Y3862 | SparPtdh3-YFP-cyc1 | Reference | Reference |
| Y3824 | ScerPtdh3-YFP-cyc1 | ∆TDH3 | Reference |
| Y3910 | SparPtdh3-YFP-cyc1 | ∆TDH3 | Reference |
| Y3908 | SmikPtdh3-YFP-cyc1 | ∆TDH3 | Reference |
| Y3909 | SkudPtdh3-YFP-cyc1 | ∆TDH3 | Reference |
| Y4128 | ScerPar5Ptdh3-YFP-cyc1 | Reference | Reference |
| Y4135 | SparCer5Ptdh3-YFP-cyc1 | Reference | Reference |
| Y4168 | ScerPtdh3-YFP-cyc1 | Reference | ∆Tye7 |
| Y4179 | SparCer5Ptdh3-YFP-cyc1 | Reference | ∆Tye7 |
| Y4180 | SparPtdh3-YFP-cyc1 | Reference | ∆Tye7 |
| Y4181 | ScerPar5Ptdh3-YFP-cyc1 | Reference | ∆Tye7 |
