## Supplementary Tables for "*cis-* and *trans*-regulatory factors contributing to divergent activity of the *TDH3* promoter in *Saccharomyces* yeast"

**Supplementary Table 1A. Genomic Background: Reference**

| Contrast | Estimate (log10-scaled) | SE | df | lower.CL | upper.CL | t.ratio | p.value |
|---|---|---|---|---|---|---|---|
| S.cer - S.par | 0.2995 | 0.057 | 18.4 | 0.139 | 0.460 | 5.266 | 0.000 |
| S.cer - S.mik | 0.2428 | 0.057 | 18.4 | 0.082 | 0.403 | 4.269 | 0.002 |
| S.cer - S.kud | 0.2279 | 0.057 | 18.4 | 0.068 | 0.388 | 4.007 | 0.004 |
| S.par - S.mik | -0.0567 | 0.058 | 18.0 | -0.221 | 0.108 | -0.973 | 0.766 |
| S.par - S.kud | -0.0716 | 0.058 | 18.0 | -0.236 | 0.093 | -1.230 | 0.617 |
| S.mik - S.kud | -0.0149 | 0.058 | 18.0 | -0.180 | 0.150 | -0.256 | 0.994 |

**Supplementary Table 1B. Genomic Background: *TDH3* Deletion**

| Contrast | Estimate (log10-scaled) | SE | df | lower.CL | upper.CL | t.ratio | p.value |
|---|---|---|---|---|---|---|---|
| S.cer - S.par | 0.4875 | 0.095 | 19.7 | 0.221 | 0.754 | 5.130 | 0.000 |
| S.cer - S.mik | 0.2988 | 0.095 | 19.7 | 0.032 | 0.565 | 3.144 | 0.025 |
| S.cer - S.kud | 0.2647 | 0.095 | 19.7 | -0.002 | 0.531 | 2.785 | 0.052 |
| S.par - S.mik | -0.1887 | 0.092 | 18.0 | -0.449 | 0.072 | -2.049 | 0.208 |
| S.par - S.kud | -0.2228 | 0.092 | 18.0 | -0.483 | 0.037 | -2.419 | 0.109 |
| S.mik - S.kud | -0.0341 | 0.092 | 18.0 | -0.294 | 0.226 | -0.371 | 0.982 |

**Supplementary Table 1.** Estimated contrasts of reporter expression values calculated using the *emmeans* package in R from a linear-model with promoter genotype, *TDH3* genotype, and the interaction between promoter genotype and *TDH3* genotype included as fixed-effect variables. The experimental day was included as a random-effect variable. Degrees-of-freedom were estimated for this mixed-effects model using the Kenward-Roger method and *P*-values were adjusted using the Tukey HSD method.

**Supplementary Table 2A. Estimated marginal means**

| Reporter | Tye7 | estimate | SE | df | Lower-CL | Upper-CL | estimate (%) | Lower-CL(%) | Upper-CL(%) |
|---|---|---|---|---|---|---|---|---|---|
| Scer_cer5nt | Tye7 | 1.0001 | 0.0048 | 52.11 | 0.9903 | 1.0098 | 100.02 | 97.80 | 102.28 |
| Scer_par5nt | Tye7 | 0.9824 | 0.0048 | 52.11 | 0.9726 | 0.9921 | 96.02 | 93.89 | 98.20 |
| Spar_cer5nt | Tye7 | 0.9581 | 0.0048 | 52.11 | 0.9484 | 0.9679 | 90.81 | 88.80 | 92.87 |
| Spar_par5nt | Tye7 | 0.9340 | 0.0048 | 52.11 | 0.9243 | 0.9437 | 85.90 | 83.99 | 87.84 |
| Scer_cer5nt | ΔTye7 | 0.9148 | 0.0048 | 52.11 | 0.9051 | 0.9246 | 82.19 | 80.37 | 84.05 |
| Scer_par5nt | ΔTye7 | 0.9020 | 0.0048 | 52.11 | 0.8923 | 0.9117 | 79.80 | 78.03 | 81.61 |
| Spar_cer5nt | ΔTye7 | 0.8905 | 0.0048 | 52.11 | 0.8808 | 0.9002 | 77.72 | 75.99 | 79.48 |
| Spar_par5nt | ΔTye7 | 0.8762 | 0.0051 | 57.12 | 0.8660 | 0.8863 | 75.19 | 73.45 | 76.97 |

**Supplementary Table 2B. Genomic Background: Reference**

| contrast | estimate | SE | df | t.ratio | p.value |
|---|---|---|---|---|---|
| Scer_cer5nt - Scer_par5nt | 0.0177 | 0.0061 | 160.0 | 2.908 | 0.0214 |
| Scer_cer5nt - Spar_cer5nt | 0.0419 | 0.0061 | 160.0 | 6.886 | <.0001 |
| Scer_cer5nt - Spar_par5nt | 0.0661 | 0.0061 | 160.0 | 10.853 | <.0001 |
| Scer_par5nt - Spar_cer5nt | 0.0242 | 0.0061 | 160.0 | 3.978 | 0.0006 |
| Scer_par5nt - Spar_par5nt | 0.0484 | 0.0061 | 160.0 | 7.945 | <.0001 |
| Spar_cer5nt - Spar_par5nt | 0.0242 | 0.0061 | 160.0 | 3.967 | 0.0006 |

**Supplementary Table 2C. Genomic Background: Tye7 Deletion**

| contrast | estimate | SE | df | t.ratio | p.value |
|---|---|---|---|---|---|
| Scer_cer5nt - Scer_par5nt | 0.0128 | 0.0061 | 160.0 | 2.106 | 0.1555 |
| Scer_cer5nt - Spar_cer5nt | 0.0243 | 0.0061 | 160.0 | 3.994 | 0.0006 |
| Scer_cer5nt - Spar_par5nt | 0.0387 | 0.0063 | 161.0 | 6.181 | <.0001 |
| Scer_par5nt - Spar_cer5nt | 0.0115 | 0.0061 | 160.0 | 1.887 | 0.2376 |
| Scer_par5nt - Spar_par5nt | 0.0258 | 0.0063 | 161.0 | 4.130 | 0.0003 |
| Spar_cer5nt - Spar_par5nt | 0.0143 | 0.0063 | 161.0 | 2.293 | 0.1039 |

**Supplementary Table 2.** Estimated contrasts of reporter expression values calculated using the *emmeans* package in R from a linear-model with promoter genotype, *TYE7* genotype, and the interaction between promoter genotype and *TYE7* genotype included as fixed-effect variables. The experimental day was included as a random-effect variable. Degrees-of-freedom were estimated for this mixed-effects model using the Kenward-Roger method and *P*-values were adjusted using the Tukey HSD method.
